# Response-time behaviors of intercellular communication network motifs

**DOI:** 10.1101/136952

**Authors:** Kevin Thurley, Lani F Wu, Steven J Altschuler

## Abstract

Cell-to-cell communication networks have critical roles in diverse organismal processes, such as coordinating tissue development or immune cell response. However, compared to intracellular signal transduction networks, the function and engineering principles of cell-to-cell communication networks are far less understood. Here, we study cell-to-cell communication networks using a framework that models the input-to-output relationship of intracellular signal transduction networks with a single function—the response-time distribution. We identify a prototypic response-time distribution—the gamma distribution—arising in both experimental data sets and mathematical models of signal-transduction pathways. We find that simple cell-to-cell communication circuits can generate bimodal response-time distributions, and can control synchronization and delay of cell-population responses independently. We apply our modeling approach to explain otherwise puzzling data on cytokine secretion onset times in T cells. Our approach can be used to predict communication network structure using experimentally accessible input-to-output measurements and without detailed knowledge of intermediate steps.

## Introduction

In multicellular organisms, cells live in communities and constantly exchange signaling molecules. Prominent examples of short-range communication are diffusible ligands shaping immune responses [1] and the tumor microenvironment [2], notch-delta mediated signals [3] and micro-vesicles [4]. In the mammalian immune system, cell-to-cell communication networks often consist of 10’s of cytokine species (small diffusive messenger proteins), several types of T cells and other immune cells like neutrophils and macrophages, and also epithelial cells [1,5]. In many cases, cytokines secreted by one cell type act on other cell types, but at the same time have effects on the original cell type as well: an important example is IFN-γ which is secreted by Th1 cells (a sub-class of T cells), stimulates macrophages and also induces the differentiation of T cells toward Th1 cells. Accordingly, the levels of various cytokine species vary by an order of magnitude or more between supernatants of isolated cells and cell populations [6,7], demonstrating pronounced effects of intercellular interaction networks on the cytokine milieu.

On the cellular level, extensive research has identified many molecules and pathways involved in signal transduction and, in many cases, has also developed an understanding of their function. In particular, the identification and analysis of generic network motifs has led to an understanding of how certain interaction topologies can function to suppress noise, amplify signals or provide robustness [8–12]. For this purpose, mathematical models of simplified systems have often been an important driving force, which have helped to reveal engineering principles such as feedback control and perfect adaptation [13–15]. However, on the level of cell-to-cell communication, the mapping from general network motif to function is poorly understood. It is unclear how well models that focus on specific cases, such as for IL-2 [16–19], IFN-γ [20,21] or TNF-α [22,23] signaling networks, can be used to infer properties of general network motifs. Further, intracellular networks—which are the building blocks of intercellular networks—may themselves be quite complex. Can we investigate behaviors of intercellular networks without requiring knowledge of detailed intracellular networks, whose parameters are often inaccessible with current experimental approaches?

Here we propose response-time modeling as a framework to unify and interpret knowledge on intracellular and intercellular signaling pathways. In this framework, the input-to-output response statistics of intracellular signaling networks are captured by a single function: the response-time distribution. This distribution, which describes the dynamics of cell-state switching, can be either measured directly or calculated from models describing the network. Importantly, focusing on response-time distributions allows us to elide detailed descriptions of intermediate intracellular signaling steps and focus on population-level behaviors that emerge from connecting cell-state outputs to each other via intercellular networks. Below, we first characterize response-time distributions that can arise from intracellular networks and find that, in many cases, these distributions and cellular output characteristics can be well modeled by a gamma distribution. Second, we use this observation to analyze common cell-to-cell communication network motifs, and discover that different interaction topologies can regulate a rich set of dynamic behaviors, including delayed, synchronized and bimodal cellular responses to a stimulus. Finally, we apply our approach to investigate a recent data set on cytokine secretion onset times.

## Results

### Response-time modeling of cell-state dynamics

Temporal heterogeneity has been widely observed for cellular processes that involve cell-to-cell communication, such as onset of cytokine secretion or cell differentiation (Figure 1A and Table 1). The distributions of times at which cellular responses occur may not be described by a simple Poisson process, which would show exponentially distributed response times (Figure 1B) [24]. Rather, single-peaked and even bimodal distributions can occur, reflecting the complex networks underling many biological processes.

**Figure 1:**
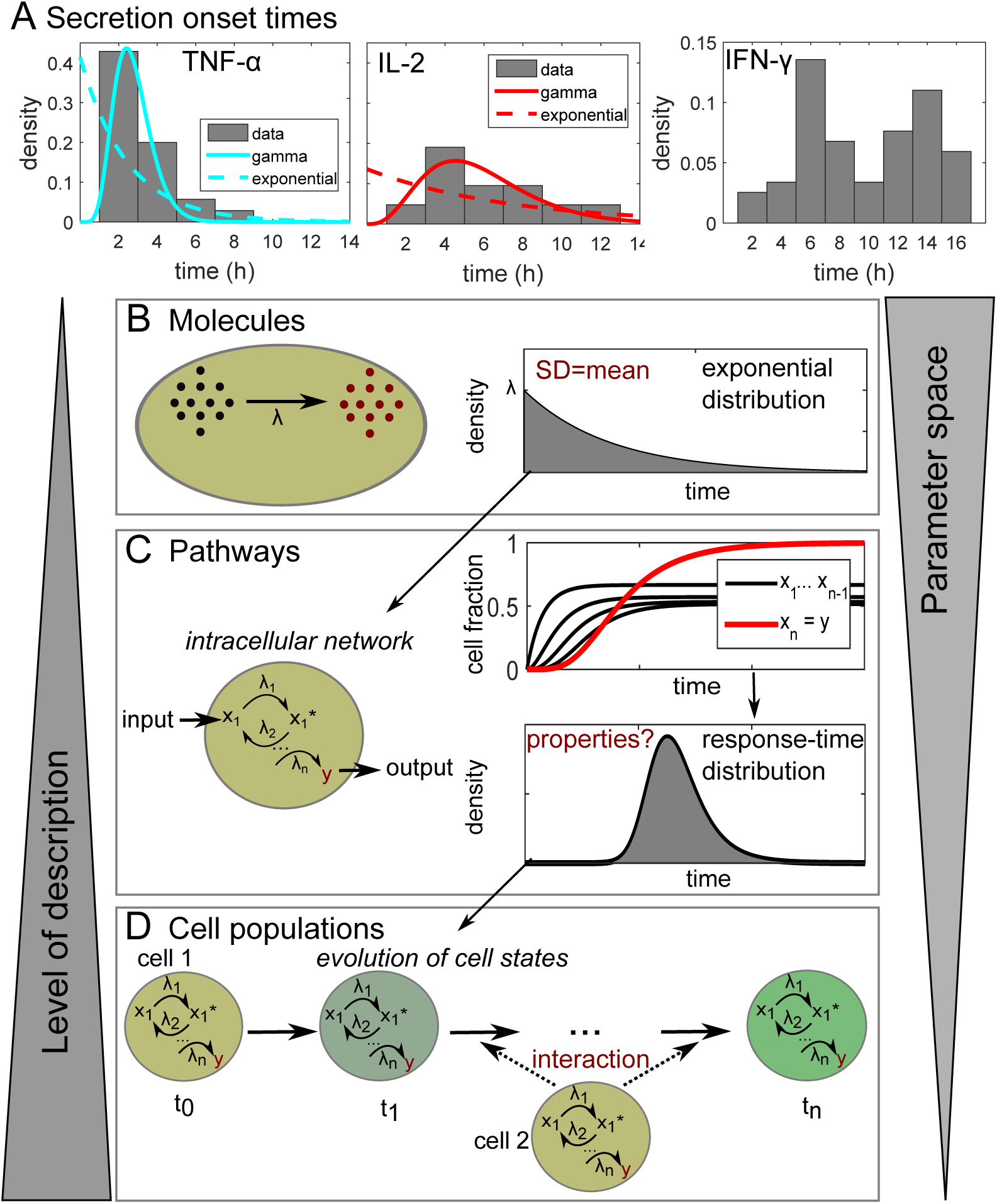
Response-time modeling of cell-state dynamics. (A) Cytokine secretion onset times in CD8+ T cells from Han et al. [34]. Data were taken from the original publication and re-normalized to probability distributions. For TNF-α and IL-2, also best-fit curves to the gamma distribution and the exponential distribution are shown (fitting parameters in Table S1). (B) An elementary chemical reaction is well described by a simple rate equation, with a single rate parameter *λ* (concentration per time). However, the lack of information on microscopic properties like positions and velocities of reacting particles implies that the waiting time until the next reaction occurs (the response time) is a random variable. Chemical reaction kinetics dictate that the response times are exponentially distributed. (C) Cellular state changes require a set of chemical reactions forming an intracellular reaction network. That network can be described by differential equations for each reaction, whose solutions reveals the fraction of cells containing each molecular species at every time point. From that information, we can calculate the response-time distribution for a cell state of interest. That response-time distribution does not need to be exponential or monotonic, but can have one or even several peaks. (D) The response of a cell population to a stimulus is often not only dependent on intracellular networks, but may also evolve by intercellular communication. Response-time modeling uses the response-time distributions for all considered cell-state changes, and their dependence on other cell states, to characterize the intercellular communication network.

**Table 1:**
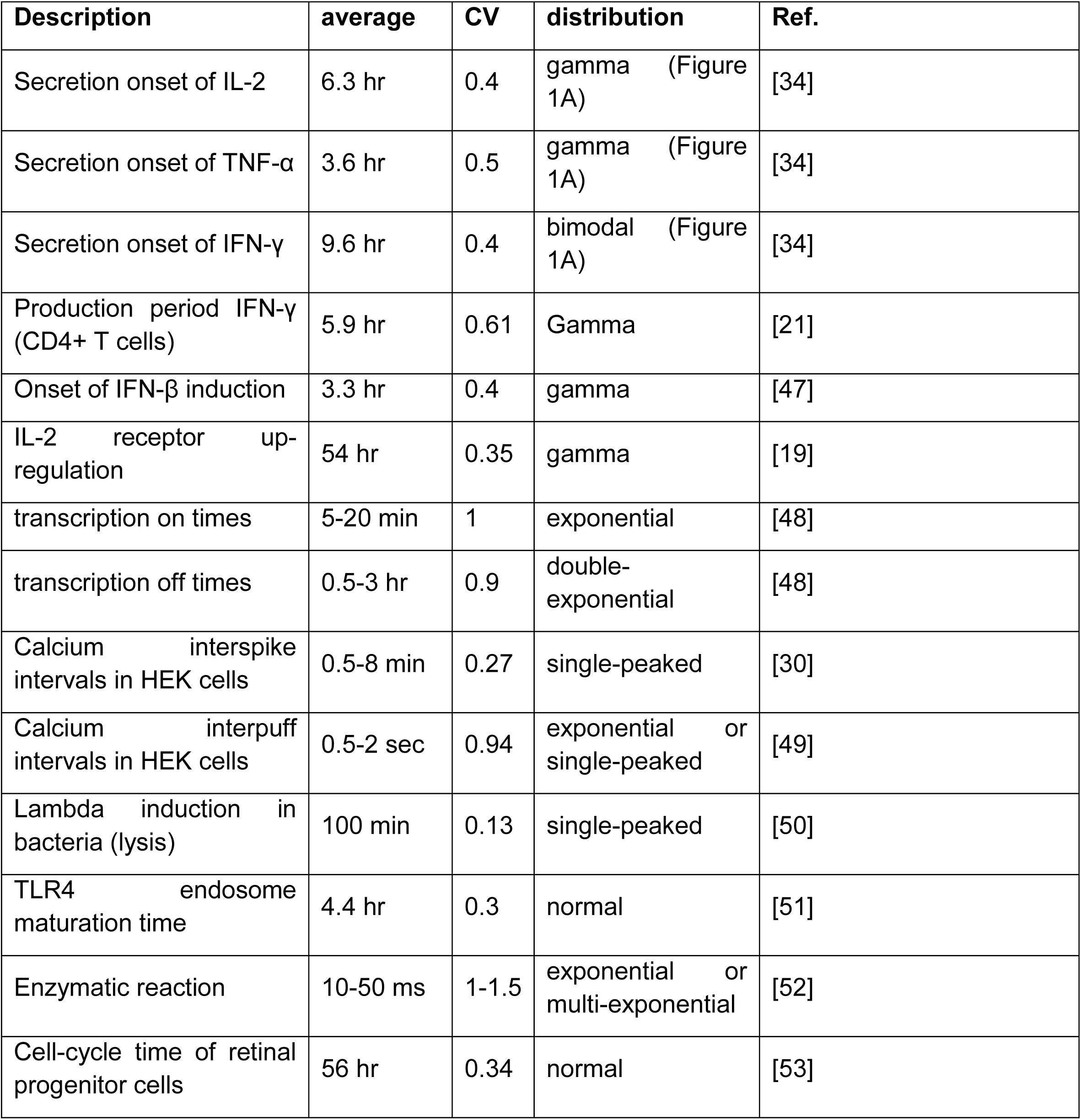
Literature survey of response-time distributions. Note that normal, gamma and double-or multi-exponential distributions all fall into the class of ‘single-peaked’ distributions. CV: Coefficient of variation.

| Description | average | CV | distribution | Ref. |
| --- | --- | --- | --- | --- |
| Secretion onset of IL-2 | 6.3 hr | 0.4 | gamma (Figure 1A) | [34] |
| Secretion onset of TNF-α | 3.6 hr | 0.5 | gamma (Figure 1A) | [34] |
| Secretion onset of IFN-γ | 9.6 hr | 0.4 | bimodal (Figure 1A) | [34] |
| Production period IFN-γ (CD4+ T cells) | 5.9 hr | 0.61 | Gamma | [21] |
| Onset of IFN-β induction | 3.3 hr | 0.4 | gamma | [47] |
| IL-2 receptor up-regulation | 54 hr | 0.35 | gamma | [19] |
| transcription on times | 5-20 min | 1 | exponential | [48] |
| transcription off times | 0.5-3 hr | 0.9 | double-exponential | [48] |
| Calcium interspike intervals in HEK cells | 0.5-8 min | 0.27 | single-peaked | [30] |
| Calcium interpuff intervals in HEK cells | 0.5-2 sec | 0.94 | exponential or single-peaked | [49] |
| Lambda induction in bacteria (lysis) | 100 min | 0.13 | single-peaked | [50] |
| TLR4 endosome maturation time | 4.4 hr | 0.3 | normal | [51] |
| Enzymatic reaction | 10-50 ms | 1-1.5 | exponential or multi-exponential | [52] |
| Cell-cycle time of retinal progenitor cells | 56 hr | 0.34 | normal | [53] |

We wondered whether cell-to-cell communication networks could be modeled and analyzed in a way that abstracts molecular detail yet still captures essential dynamic properties. For example, to fully describe an elementary chemical reaction, one would need to know the positions and velocities of all molecules at all times [24]. However, the process can be well approximated by a single phenomenological parameter, the reaction rate-constant. To develop an analogous approach for modeling cellular state changes, one must take into account that cell-state changes are the consequence of intracellular multi-step processes: the response of a cell to an input signal is not a single-step reaction but rather a result of a reaction network (Figure 1C). The time until this observable state change happens is a random variable, just like the time until the next molecular event in a single-step reaction (Figure 1B and C)—however, it is in general not exponentially distributed. Rather, this response-time distribution is a signature function depending on and describing the relevant intracellular processes, with no known *a priori* properties.

Response-time modeling has the advantage that we do not need to take all details of intracellular dynamics into account, but rather focus on the key measurable events (see also [25,26]). Therefore, compared to models on the level of molecular species or even individual molecules, we can describe the behaviors of cell populations with a rather small number of parameters (few measurable response-time distributions instead of 100’s of poorly accessible rate parameters) (Figure 1D). To assess what insights response-time modeling can give us on the dynamics of cell-state networks, the first step was a characterization of response-time distributions resulting from simple intracellular multi-step processes—the building blocks of more complex networks.

We note that throughout the paper, we refer to multi-step processes as “intracellular networks” and to their superposition as “cell-to-cell signaling”. In fact, in a mathematical sense, all these processes can be viewed as components of one large network, and in many cases, it is somewhat arbitrary which parts of that network are isolated and considered as “subnetworks” that are subsumed by a response-time distribution: In other circumstances, compartments inside a cell might be regarded as interacting subunits, or several cells in a certain micro-environment might form a community that interacts with other communities—for example via hormones or neural circuits. All those cases can, in principle, be treated analogously to the framework of response-time modeling presented here.

### Simple intracellular networks induce single-peaked response-time distributions

In a literature survey, we found that many reports of experimentally measured response-time distributions indicate a single-peaked type of distribution. Such distributions have been reported for a wide range of cellular systems from gene transcription over cellular Ca^2+^ spikes to cytokine secretion (Table 1 and Figure 1A), with the exception of some processes where exponential distributions have been measured, and bimodal IFN-γ secretion onset times in T cells, which we discuss in detail later.

Why does this widespread occurrence of single-peaked response-time distributions occur, and what does it mean for the typical dynamics of a cell population? Response times for single-step reactions are exponentially distributed (Figure 1B and Figure 2A, top). However, cellular signal transduction typically is driven by intracellular networks comprising phosphorylation cascades, feedback, crosstalk etc. As a simple illustration, consider a uniform, irreversible reaction chain, i.e. the cellular response is triggered after completion of *n* reaction steps all driven by the same rate constant *μ* = λ/n (Figure 2B, top). This process has the same average response time as a single reaction with rate λ, but the distribution of the response times over a cell population changes: The process can be regarded as a sum of *n* single-step processes (elementary reactions), and therefore the over-all response time is the *n-*fold convolution [27] (see SI Text)

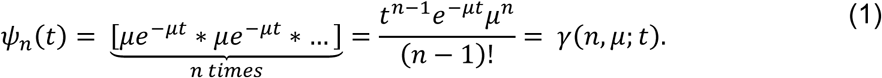

**Figure 2:**
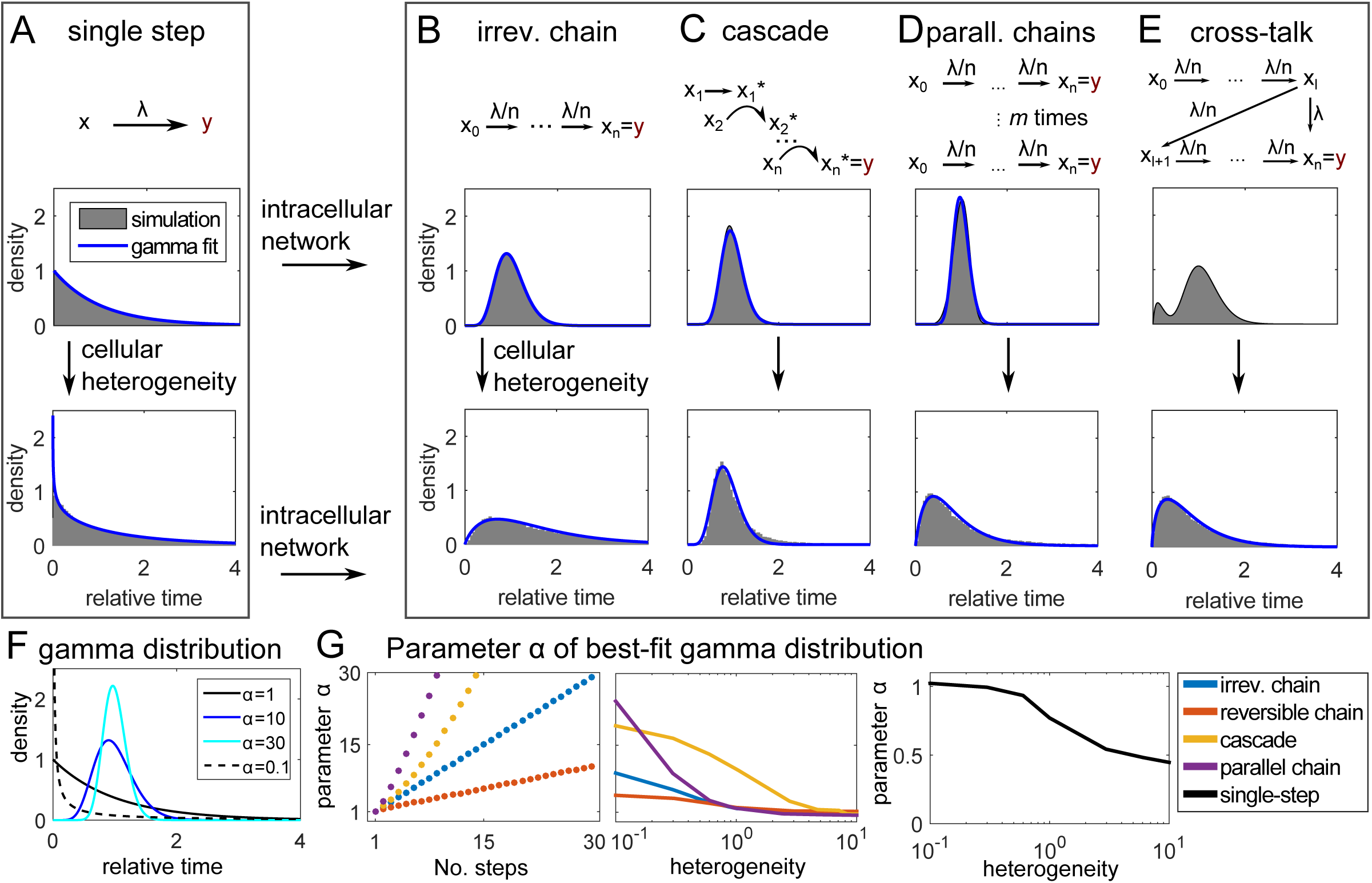
Intracellular reaction networks are often well described by gamma-distributed response times. (A-E) Response-time distributions of multi-step models. (top): In the shown models, each arrow represents an elementary (i.e. single-step) reaction (see SI Text for model equations). Response-time distributions are computed by solving the corresponding system of differential equations and normalizing by the distribution average. (bottom): To account for cellular heterogeneity, the rate parameter λ is drawn from a log-normal distribution (standard deviation=mean), and normalized response-time distributions are obtained by stochastic simulation. For all models, heterogeneous λ results in longer tails and earlier peaks. Blue lines: Best-fit gamma distributions. Parameters: *n*=10, *λ*=1, *l*=1. (F) Plots of the gamma distribution (Equation 1) with rate parameter β=1/α (i.e. the average time is constant) and shape parameter as indicated. (G) Shape parameter *α* of best-fit gamma distributions to the indicated models (panel A-D and Figure S1A). “No. steps”: Parameter *n* in the models. Cellular heterogeneity: Coefficient of variation of the log-normal distribution generating *λ*.

Here, ‘*’ denotes convolution and *γ*(α, β; *t*) is known as the gamma distribution with shape parameter *α* and rate parameter *β* (in general, α can take non-integer values, see SI Text).

Indeed, the single-peaked response-time distributions observed experimentally can be described by gamma distributions (Table 1), as for α>1, the gamma-distribution is an asymmetric (right-skewed) distribution with a single peak at *t*>0. The observed exponential distributions for single-enzyme kinetics, offset of transcription and intracellular Ca^2+^ puffs, indicate single-step processes (Figure 2A): All these processes are likely dominated by a single molecular reaction (binding of a metabolite to an enzyme, unbinding of a promoter from DNA, opening of a Ca^2+^ channel subunit).

Intracellular signaling pathways are usually not simple irreversible chains, and therefore, we asked whether the observed single-peaked distributions can be generated by a broader class of intracellular network models. Indeed, single-peaked distributions have previously been reported for more realistic models of cellular signal transduction like kinetic proofreading [28], multiple phosphorylation [29] and Ca^2+^ signaling [30]. Here, we studied three additional simple network motifs in more detail: The signaling cascade [8], a set of parallel irreversible chains reflecting *m* receptor molecules that each can trigger a cellular response as a “race to the nucleus” [29] (Figure 2C-D, top), and the reversible chain (Figure S1A). The response times of all those examples are well approximated by gamma distributions (Figure 2C-D and S1A, blue fitting lines, and Figure S1B and C).

Apart from intracellular networks, another complication is cellular heterogeneity: In a cell population, even a clonal one, we cannot expect that each cell has the same reaction rate for a certain intracellular process. Rather, gene expression and receptor expression levels show heterogeneity [31]. To investigate the effect of such heterogeneity on the response-time distribution, we used log-normal distributed reaction-rate parameters (Figure 2A-E and Figure S1A, bottom). In all models, the response-time distribution shifts towards longer tails and earlier peaks after incorporating cellular heterogeneity, but is still well approximated by a gamma distribution (Figure 2A-D, blue fitting lines, and Figure S1B). This is expected: Intuitively, adding cellular heterogeneity should reduce predictability; however, adding high-numbers of intermediate intracellular steps increases the predictability of the process due to the central limit theorem, which tightens the peak in the response time (Figure 2F-G)[27].

Thus far, we only studied unbranched multi-step processes. A final interesting case is cross-talk within an intracellular multi-step process (Figure 2E). In this case, a bimodal response-time distribution can occur, but even here, heterogeneity of rate parameters shifts the distribution towards a gamma-distribution (Figure 2E, bottom panel), offering another explanation for the versatility of gamma distributions. Therefore, in the following discussion, we will focus on cell population responses that induce gamma-distributed response times.

### Response-time modeling of intercellular network motifs

Having established the typical response-time patterns emerging from intracellular processes, we next asked how more general cell-state transitions shape dynamic response patterns of cell populations. For this purpose, we made use of response-time modeling (Figure 3A), which describes a cell-state change from a state *S*_*i*_ to *S*_*j*_ by a semi-Markov process defined by response-time distributions *ψ*_*ij*_(*t* – *τ*) (also known as first-passage times; see SI Text for precise definitions). Here we specifically chose gamma-distributed response times (Equation 1), because of their frequent occurrence in intracellular processes (Table 1 and Figure 2B-E). An advantage of this approach is that we can consider cell-to-cell interactions including feedback (e.g. by exchange of diffusible ligands) simply as a dependence of the parameters of the gamma distribution on the fraction of cells in a certain cellular state *S*_*l*_:

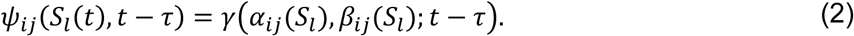

**Figure 3:**
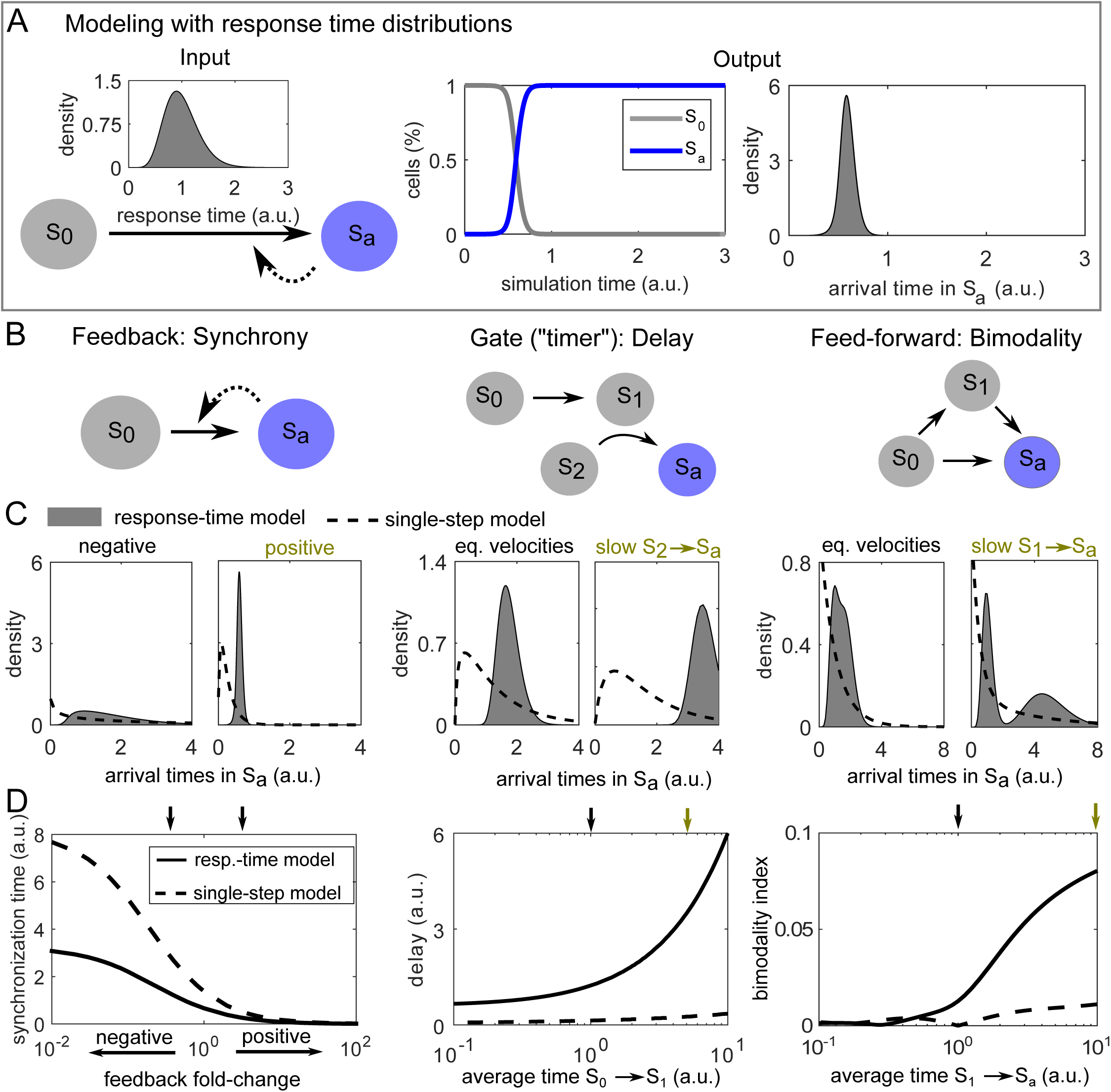
Network motif analysis using response-time modeling. (A) Illustration of response-time modeling: Each reaction arrow represents an intracellular multi-step process represented by a gamma-distribution *γ*(α, β; *t*) (Equation 1). The process is started in state *S*_*0*_ and continues until all cells reach the absorbing state *S*_*a*_. Dashed arrow: Positive feedback. Arrival time: *A posteriori* distribution of the times to reach state *S*_*a*_ considering feedback. (B-D) Simple models (network motifs) of cell-to-cell communication. To keep the response-time models and single-step models comparable, we scaled the rate parameter of the gamma distributions as β → αβ, so that the average of the distribution is 1/β independently of α. Feedback and interaction (gate motif) are modeled by Michaelis-Menten type equations (see Methods). Parameter values used in (C) are indicated by small color-coded arrows in (D). ‘Average time’: Average 1/β^base^ of the gamma distribution representing the respective reaction. Parameters not stated otherwise: *α*=10 (“response-time model”) or *α*=1 (“single-step model”, i.e. the exponential distribution is used), *K*=0.1, β^base^=1, feedback fold-change η=5 (positive feedback) and 0.2 (negative feedback). In the feed-forward loop motif, the branching probability is *p*_01_ = *p*_02_ = 0.5.

To completely determine the system, one needs to also provide probabilities *p*_*ij*_ for the execution of each possible reaction (with ∑_*i*_ *p*_*ij*_ = 1), e.g. in the case of branching reactions. Note that in the basic framework presented here (Equation 2), we assume a “well-stirred” situation and do not take into account spatial effects like concentration gradients in diffusible messengers [17].

Using response-time modeling, we analyzed a set of simple toy models or “network motifs” that often appear in larger cell-to-cell communication networks (Figure 3B). In analogy to the simple multi-step models studied in Figure 2, our main readout is the *a posteriori* distribution of arrival times in an absorbing state *S*_*a*_ (Figure 3C) (in general, every state of interest can be treated as an absorbing state by removing transitions leaving that state [27]). In contrast to the response-time distributions in Figure 2, the arrival-time distributions are not normalized, but are analyzed separately within each motif to reveal the effect of parameter values on the delay and synchronization time and on bimodality (Figure 3D)(see Methods).

As a first example, consider a single cell-state transition with feedback (Figure 3B-D, “feedback”). We found that positive feedback decreases and negative feedback increases the width of the arrival-time distribution. To quantify this property, we defined the “synchronization time” as the minimal time frame in which a certain fraction of cells (here 75%) responds after an initial delay time (see Methods). Feedback has only minor effects on this delay time (Figure S2A). Thus, feedback regulation between cells is well suited to generate highly synchronized or desynchronized responses across a cell population. Conversely, we asked whether a simple cellular communication network could control the delay without changing synchronization—a sort of “timer” circuit for the cellular population. Indeed, we found that long delays can be achieved without increased synchronization times by adding a bottleneck, e.g. in terms of the positive interaction or “gate” motif (Figure 3B-D, “gate”). The gate motif increases delay without changing synchronization over a wide parameter range (Figure S2A). Adding delay by simply slowing down the intracellular processes that induce one or two consecutive cell-state changes is not sufficient for this effect, as here the synchronization time increases substantially when adding delay (Figure S2B-D). Intuitively, the higher level of synchronization in the gate motif can be explained by the global positive interaction, which increases synchronization similar to the positive feedback case, and therefore compensates for the loss of synchronization due to a shorter time scale.

Finally, we studied the redundant, coherent feed-forward loop. This motif is a simple model of the situation that cellular activation (reaching state *S*_*a*_) can be induced in several different ways, for example by means of different types of cytokines. Quite interestingly, we found that this motif can generate a bimodal distribution of arrival times in the absorbing state (Figure 3B-D, “feed-forward”). Intuitively, that bimodality is caused by the contributions from the “direct” and the “indirect” (via *S*_*1*_) ways to reach *S*_*a*_. However, substantial bimodality only arises if there is a time-scale separation between the two routes to *S*_*a*_, as implemented here by a longer average time in the process *S*_1_ → *S*_*a*_; otherwise there is no clear separation between the two peaks, giving rise to a single long-lasting cellular response of moderate intensity (i.e. larger synchronization time, see Figure S2A). While bimodal distributions can also occur by crosstalk inside intracellular networks (essentially also a feed-forward loop)(Figure 2E), a feed-forward loop of elementary reactions is not sufficient for bimodality (Figure 3B-D, dashed lines): In general, our analysis suggests that at least three consecutive elementary steps are required for a bimodal response-time distribution (SI Text).

### Cell-to-cell communication allows independent control of delay and synchronization

Above, we used the prototypic response-time distributions arising in intracellular signal transduction networks to analyze common network motifs of cell-state dynamics. We found that in the framework of response-time modeling, simple network motifs can induce emergent behavior such as bimodal response times, which does not arise in the corresponding single-step model that neglects the multi-step nature of cell-state changes. Moreover, we found that circuits, such as the feedback or gate motif, can control synchronization and delay independently (Figure 4). Our simulations strongly indicate that independent control is not possible with intracellular multi-step processes alone (Figure 4, blue line): in processes that can be described by a gamma distribution, synchronization and delay are closely related by a linear relation.

**Figure 4:**
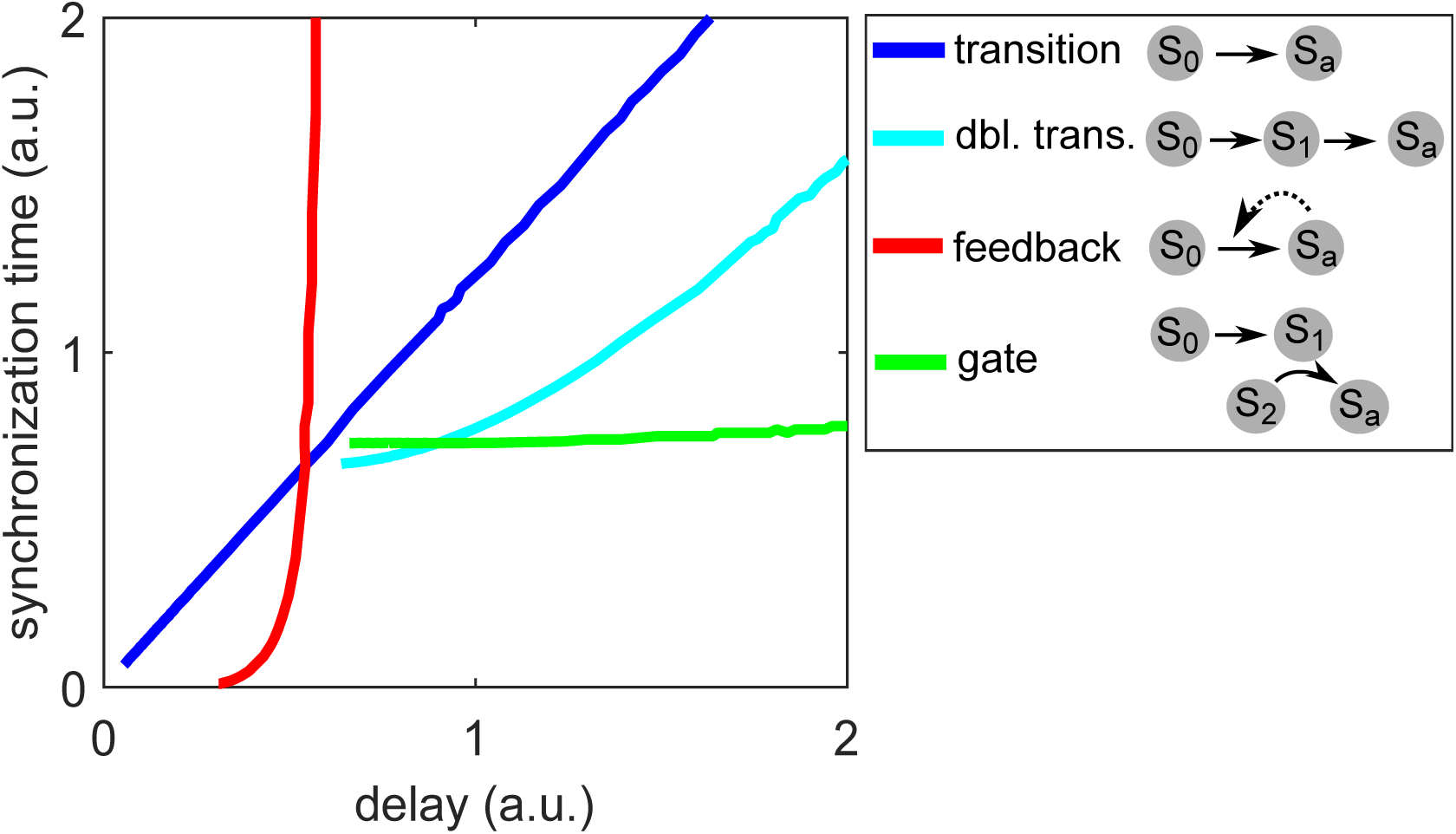
Independent control of delay and synchronization. Comparison of the gate and feedback motifs (Figure 3B-D), and the single or double state transition motifs (Figure S2B-D). The gate motif allows for variable delays for the same degree of synchronization while, conversely, the feedback motif allows variable synchronization for the same delay. Synchronization and delay cannot be decoupled for the single transition, representing intracellular multi-step processes alone (see Figure 2). All curves are generated by changing the time-scale or feedback strength parameters as in Figure 3D or Figure S2D.

### Delay-induced persistence detection

A network that rejects transient activation signals and only responds to persistent signals has been termed a “persistence detector” [9,32]. Persistence detection in cell-to-cell communication has recently been demonstrated in the context of a paracrine signal induced by opto-genetic tools, which can precisely control the timing of an input stimulus [33]. The precise mechanism of persistence detection is still to be resolved, however we wondered whether it could be explained by the delay caused by early cytokine secretion onset (Figure 5A). In this model, the cell-state change of a silent “receiver cell” to a cell with detectable fluorescence signal is mediated by a paracrine cytokine signal, which is again triggered by the (time-controlled) opto-genetic stimulus. As mentioned before, cytokine-secretion onset times are often gamma-distributed (Table 1).

**Figure 5:**
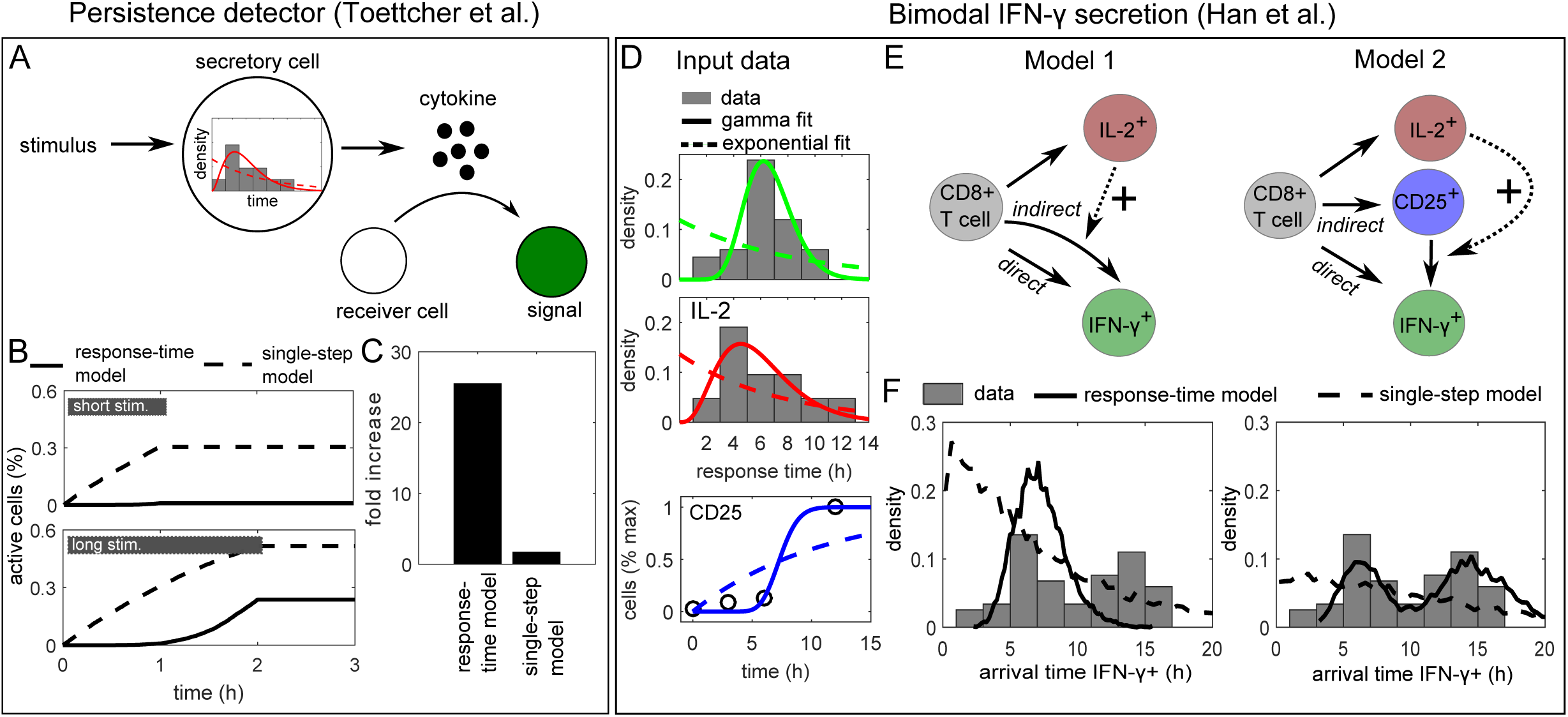
Persistence detection and bimodal IFN-γ secretion onset. (A) Persistence detection: The experiment by [33] was simulated with a stimulus ON-time of either 1hr or 2hr, triggering a cell-state change that is governed by the response-time distribution measured for secretion onset of the early cytokine IL-2 (Figure 1A). In the original experiment, receiver cells show a fluorescent signal (a state-change) when exposed to an opto-genetic stimulus for 2 hr but not after 1 hr stimulation, and a paracrine signal was essential for the effect. (B-C) Simulation of the model in (A) with the best-fit gamma distribution (“response-time model) or the best-fit exponential distribution (“single-step model). “Fold change”: Relative increase in maximal activity from short to long stimulus. (D) IFN-γ secretion onset: Input data used for the models (see also Table S1). IL-2 and IFN-γ secretion onset times were taken from [34] (Figure 1A), and the initial IFN-γ secretion onset times were obtained by cutting after the dip at 10 hr and renormalizing. Kinetics of CD25 (α-subunit of IL-2R) up-regulation were taken from [38] and normalized to maximal expression. Fitting lines show best-fit curves to gamma and exponential distributions (for CD25, the corresponding cumulative distribution function was used). (E) Response-time models of IL-2 and IFN-γ secretion onset. Solid arrows represent an intracellular multi-step process represented by a response-time distribution (Figure S5) and a probability to execute each of the branching reactions (see Table S1). Dashed arrows represent positive interaction. (F) Simulation of the models in (E): *A posteriori* arrival times to reach state “IFN-γ+”, i.e. initiate IFN-γ secretion. Note that the probabilities *p*_*ij*_ for each differentiation path (IL-2+, CD25+, IFN-g+) are determined independently of the response-time distributions (see Table S1), and therefore the shown arrival times are normalized to the corresponding fraction of cells: Given a certain differentiation path, it is certain that the new state will be reached eventually. “Response-time model”: Simulations with best-fit gamma distributions (here non-integer valued shape parameters are possible); “single-step model”: Simulations with best-fit exponential distributions.

Here, we studied a simple model where cell-state transitions are executed with the response-time distribution measured for IL-2, and are triggered by an external stimulus that is present for either 1 hr or two hr as in [33]. Indeed, we found that the response-time model, but not the single-step model, yields a strong difference in cell activation between the 1 hr and 2 hr stimuli (Figure 5B and C), providing a plausible explanation for the reported persistence detection. A more generic analysis revealed the benefit of signaling circuits that control delay independently of synchronization (Figure 4). Again, we assumed an input stimulus of limited duration (Figure S3A), now acting on a cell-state change controlled by an underlying “delay-inducing” gate or transition (Figure 4) network motif. We modeled these motifs using our response-time approach (i.e. fit gamma distributions to their input-to-output relationship) and scaled the average response times for both motifs so that they have the same delay (3 time units). Our simulations show that both delay-inducing motifs exhibit some degree of persistence detection compared with a simple, single-step process (Figure S3B), but only the gate motif allows for 100% of the cells to be activated for a long stimulus (e.g. 5 time units) while still rejecting a short stimulus (e.g. 3 time units). Moreover, the gate motif has a sharp transition in signal amplitude (Figure S3C).

### Bimodal IFN-γ secretion onset times

In our literature survey (Table 1), a striking example of a response-time distribution that deviates from the commonly observed single-peaked pattern is the bimodal IFN-γ secretion onset times [34](Figure 1A). Our analysis of intercellular communication networks suggested that a feed-forward loop motif can evoke a bimodal response-time distribution (Figure 3B-D, feed-forward). As it is known that IL-2 stimulates IFN-γ secretion of CD8+ T cells [35,36], we next examined whether a combination of direct (antigen driven) and indirect (IL-2 mediated) stimulation of IFN-γ secretion is sufficient to explain the bi-modal distribution.

Response-time modeling allows annotating cell-state models by directly using measured transition probabilities and response-time distributions. That way, we were able to completely specify the process (except for the IL-2 interaction strength) based on a published data-set [34] (Figure 5D and Table S1): The onset times of IL-2 secretion are well described by a gamma-distribution, and the same is true for the early IFN-γ onset times. For late (indirect) IFN-γ secretion, we used the same distribution modified by IL-2 interaction (Figure 5E, Model 1)(Methods). The reasoning was that likely similar pathways are involved in the production and secretion of IFN-γ in both the direct and indirect case, only that they are activated either directly by antigen or indirectly via IL-2 (possibly after weak antigenic pre-stimulation). To simulate the process, we used a generalized Gillespie algorithm [37], which is necessary here because some of the input gamma distributions have a small non-integer valued shape parameter (see Methods).

Clearly, the response-time distribution generated by Model 1 is not bi-modal, and does not explain the data even qualitatively (Figure 5F). The reason is that the initial onset time distributions for IL-2 and IFN-γ are too similar, and therefore their combination leads to a single broad peak rather than a second peak in the response times (cf. Figure 3C, feed-forward loop). Thus, we reasoned that another mechanism must account for this observed delay. In fact, unstimulated T cells express only very limited amounts of the high-affinity IL-2 receptor CD25, and therefore we asked whether stimulation-induced CD25 up-regulation may cause that additional delay (Figure 5E, “Model 2”). For this process, we used CD25 expression kinetics of CD8+ T cells measured in Ref. [38], which are also well described by a gamma distribution (Figure 5D). Indeed, model 2 generates a bimodal distribution for IFN-γ secretion onset, and is in good qualitative agreement with the reported values (Figure 5F). The corresponding single-step model (dashed line) cannot reproduce the bimodal shape of the distribution. That demonstrates that our approach using the full response-time distribution is necessary to explain the data, and cannot be replaced by single-step models using only average response times.

## Materials and Methods

### Simple Models

The model schemes in Figure 2B-E were translated into differential rate equations by standard methods (see SI Text for model equations). In all models with a single absorbing state *x*_*n*_ (all except the parallel chain) and without cellular heterogeneity in the reaction rate parameter, the response-time distribution can be obtained directly from the differential equation solutions as a first-passage time [27], 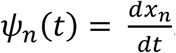. For the parallel chain parallel chain, the probability to reach the final state *n* in the first-out-of-*m* parallel processes, 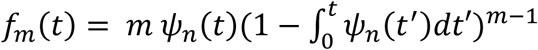 [29], is taken as the response-time distribution. To account for cellular heterogeneity, we also considered a log-normal distributed rate parameter λ. In that case, the response-time distribution is obtained by stochastic simulation (n=20000) using Gillespie’s algorithm.

### Cellular state-transition models

In general, models with several reaction channels that show non-exponential response-time distributions lead to non-linear integral equations (SI Text), which is a numerically hard problem. Instead, we employed two different methods to simulate the process: (i) Linear chain trick (used in Figures 3 and S2): If the response-time distributions are well approximated by gamma distributions with integer valued shape parameter (Equation 1), then one can replace each distribution by the corresponding irreversible *n*-step process, reducing the problem to an ODE system (see SI Text). (ii) Generalized Gillespie algorithm (Figure 5): This recently developed method [37] exploits some approximations valid for large numbers of responding cells to efficiently simulate the process with arbitrary input distributions (i.e. also gamma distributions with non-integer valued shape parameter can be used).

### Models with feedback and interaction

Feedback and interaction are modelled by a dependence of the rate parameter *β* of the input gamma distribution (Equation 1) to the fraction of cells in a state *S*_*l*_. For positive and negative feedback (Figure 3B-D, “feedback”), we used 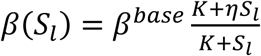 in Equation 2, where *β*^*base*^ is the base-level rate parameter, and the fold-change *η* determines feedback type and strength (positive feedback: *η* > 1, negative feedback: *η* < 1). For cellular interaction (Figure 3B-D, “gate”, and Figure 5E), we use 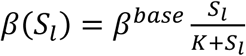.

### Measures of response-time distributions

#### Delay

We defined the delay time *t*_*delay*_ as the longest time before ≤ 5% of a cell population reach the active state, so it is the 5-percentile of the response-time distribution.

#### Bimodality

To quantify bimodality, we used the standard error (root-mean square of the sum of residuals) of a best-fit to the gamma distribution, with the rational that a bimodal distribution cannot be fit by a single gamma distribution. This approach has been widely used with normal distributions (“dip-test” [39]).

#### Synchronization

We characterized synchronization using the probability that an event occurs in (*t*, *τ*) but has not occurred before, the future life time 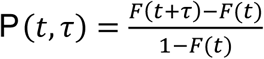, where *F(t)* is the cumulative probability distribution to the analyzed arrival-time distribution. The condition *P*(*t*, *τ*) = *d*, i.e. a fraction *d* of cells (here *d*= 75%) responds in (*t*, *τ*), together with the delay time *t*_*delay*_, defines a “synchronization time” *t*_*sync*_ = *τ*(*t*_*delay*_).

## Discussion

Response-time distributions have been explored earlier for model reduction techniques [25,26] and to analyze specific biological systems [21,23,40–42], applications including gene expression dynamics, viral infection and noise propagation in signaling pathways. Those studies demonstrate that many complex biological systems cannot be adequately described by single-step rate equation models, or at least not in their physiological environment. The reason is not only that biological networks are incompletely understood, so that known pathway maps are incomplete – rather, the fundamental problem is that the parameters needed to describe cellular networks (e.g. reaction rate parameters for all sub-processes involved in expression of a gene) cannot all be determined in vivo. Using the response-time distributions of “mesoscopic”, i.e. measurable processes, circumvents that problem by abstracting from the underlying microscopic dynamics, in analogy to chemical reaction kinetics (Figure 1). In case of branching reactions, we additionally assign probabilities for taking each path (Figures 3B and 5E). That approach is more general than assuming stochastic competition [43,44] (meaning that the earliest event wins), and branching probabilities are often available from end-point data such as flow-cytometry measurements.

The analysis of common network motifs has a long tradition in systems biology, and was used to elucidate metabolic networks [45] and gene regulatory networks [9,11,15], amongst others. The reasoning is that large, physiologic networks are composed of small, functional network motifs and can be rationalized based on these building blocks. To demonstrate such an approach for cell-to-cell signaling, we elucidated two published examples of intercellular interaction. We found: (i) that the paracrine persistence detector [33] can be explained by a delayed response-time distribution, which possibly stems from the onset of cytokine secretion; and (ii) our analysis of IFN-γ secretion onset times [34] revealed that the observed secondary response can be explained by a feed-forward loop motif consisting of IL-2 secretion and IL-2 receptor up-regulation. Those results provide a rationale for plausible mechanisms that can be tested experimentally in future research. Moreover, both examples demonstrate an advantage of our modeling approach, which is that no or very few free parameters need to be assigned if the response-time distributions of key processes are measured directly.

Cell-to-cell interaction is crucial for many functions of higher organisms, and complex intercellular communication networks have been discovered over the last decades. While the experimental capabilities to elucidate cellular responses to specific input stimuli are becoming increasingly available—sometimes even for spatiotemporal, single-cell analysis [46]—there will always be missing information. Response-time modeling offers a timely approach for predicting communication network structure and behavior using experimentally accessible input-to-output measurements even without detailed knowledge of intermediate steps.

## Acknowledgments

We would like to thank Sigurd Angenent, Orion Weiner, Michael Chevalier and members of the Altschuler and Wu lab for helpful discussions and critical feedback. This work was supported by a Research Fellowship from the German Research Association DFG (K.T.), the National Institute of Health grants GM112690 (S.J.A.) and R01CA185404 (L.F.W.), the UCSF Program for Breakthrough Biomedical Research (S.J.A., L.F.W.), and the Institute of Computational Health Sciences (ICHS) at UCSF (S.J.A., L.F.W.).

